# Cell-context dependent in silico organelle localization in label-free microscopy images

**DOI:** 10.1101/2024.11.10.622841

**Authors:** Nitsan Elmalam, Assaf Zaritsky

## Abstract

In silico labeling prediction of organelle fluorescence from label-free microscopy images has the potential to revolutionize our understanding of cells as integrated complex systems. However, out-of-distribution data caused by changes in the intracellular organization across cell types, cellular processes or perturbations, can lead to altered label-free images and impaired in silico labeling. We demonstrated that incorporating biological meaningful cell contexts, via a context-dependent model that we call CELTIC, enhanced in silico labeling prediction and enabled downstream analysis of out-of-distribution data such as cells undergoing mitosis, and cells located at the edge of the colony. These results suggest a link between cell context and intracellular organization. Using CELTIC to generate single cell images transitioning between different contexts enabled us to overcome inter-cell variability toward integrated characterization of organelles’ alterations in cellular organization. The explicit inclusion of context has the potential to harmonize multiple datasets, paving the way for generalized in silico labeling foundation models.

## Introduction

Organelles act in concert to shape and enable cell function. Accordingly, the organization of organelles and the spatial relations between different organelles are remarkably versatile and can change in response to multitude factors including undergoing cellular processes such as proliferation ^1^, migration ^2^ or differentiation ^3,4^, and being influenced by extrinsic factors such as local cell density and different microenvironmental conditions (e.g., mechanical stresses, diffusible factors, chemical treatments) ^5,6^. For instance, during mitosis, the nuclear envelope disassembles, the nucleus undergoes condensation and separation, the Golgi apparatus is disassembled and then reformed, and microtubules rearrange to form the mitotic spindle ^1^. The ability to measure whether and how the intracellular organization of organelles change is fundamental to cell biology but is technically challenging due to substantial limitations in simultaneous labeling of multiple organelles within the same cell ^7^.

In silico labeling of organelles is the computational cross-modality translation of label-free transmitted light microscopy images to their corresponding organelle-specific fluorescent images ^8^. In silico labeling holds the promise of enabling computationally multiplexed live cell imaging toward an integrated understanding of the cell ^9^. Attaining an in silico labeling model involves the acquisition of matched label-free and fluorescently labeled images, and using them to train a deep learning model that maps the label-free images to their corresponding matched fluorescence images ^10^. This training process is repeated for each organelle, producing a set of organelle-specific in silico labeling models (Fig. 1A, left). At inference, the organelle-specific models can be applied to generate a multiplexed image displaying the localization of several organelles simultaneously^11^ (Fig. 1A, right). Several recent studies demonstrated that in silico labeling can be applied to reveal how the intracellular organelle organization, cell shape and/or cell behavior alters in response to different cell states and different experimental conditions ^11–26^. This forthcoming wave of in silico labeling applications to cell phenotyping raises a major question regarding generalization: are in silico labeling models confounded by cells that are not sufficiently represented during training? In principle, changes in the intracellular organization can alter the cell’s optical properties, inducing out-of-distribution label-free images and impaired in silico labeling. For example, alterations in the cell’s internal organization due to changes in cell-cell adhesions in densely packed microenvironments may lead to changes in the cell’s optical properties that in turn can hamper high quality in silico labeling. Indeed, a few studies demonstrated inferior performance upon perturbations ^15^ or upon inference on a cell type different from the one seen on training ^27^. This deteriorated accuracy in organelle localization is posing a limitation in generalizing in silico labeling and hampering the possibility of using in silico labeling to understand how intracellular organelle organization is changing across cell types, throughout cellular processes and following perturbations.

**Figure 1.**
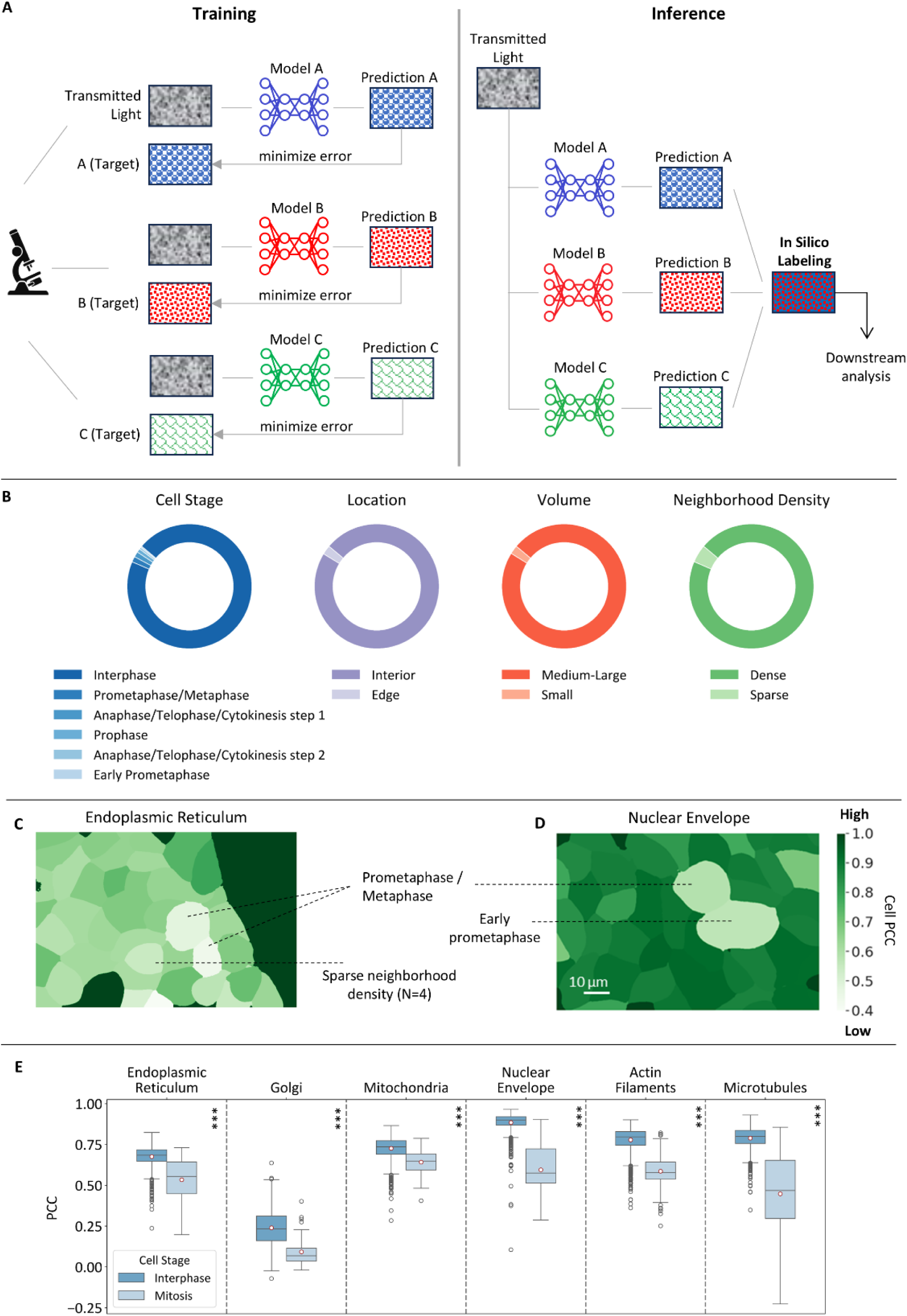
Inferior in silico labeling for rare cell populations. **(A)** In silico labeling. Training (left): an organelle-specific model receives label-free transmitted light images and their corresponding fluorescent targets and is trained to minimize the reconstruction error between the model’s prediction and the target. Inference (right): each organelle-specific model translates a transmitted light image to its corresponding predicted fluorescence image. The predictions can be combined to an integrated multi-organelle image, which can be used for downstream analyses. **(B)** Distribution of rare cell populations in the dataset, comprising 7,622 single cells. Left to right: Stage: 5% of the cells were in one of five non-interphase mitotic stages; Location: 2.4% of the cells were located at the colony edge; Volume: 2.2% of the cells had z-score lower than -1.5 relative to the population distribution; Neighborhood density: 4.7% of the cells were in sparse neighborhoods consisting of 4 or less adjacent cells. **(C and D)** Cell-level predictions. Each colored region in the field of view represents the replicated U-Net’s ^10^ average Pearson correlation coefficient of a cell. Scale bar = 10μm. (C) Poor endoplasmic reticulum in silico labeling of two cells. One cell was in the prometaphase/metaphase stage of mitosis and the other cell was in a sparse neighborhood. (D) Poor nuclear envelope in silico labeling of two cells. One cell was in the prometaphase/metaphase and the other in the early prometaphase stages of mitosis. **(E)** Distribution of single cell in silico labeling performance across organelles for cells in interphase (dark blue) versus mitosis (light blue). Mann-Whitney U test *** - p-values < 0.001. Full results including in silico labeling performance, statistical tests and population sizes are available in Table S2.

Here we focus on the problem of out-of-distribution label-free images due to rare cellular states and contexts that are under-represented in the training data. We report a decreased performance of in silico labeling for rare cell populations, and introduce a new method called *CEll in silico Labeling using Tabular Input Context*, or *CELTIC*, to overcome this limitation. CELTIC is designed to improve in silico labeling of out-of-distribution cell populations by incorporating biological meaningful cell context (encoded as tabular data) to the in silico labeling model. We show that by inclusion of cell context CELTIC enhances the in silico labeling of rare cell populations, especially organelle localization patterns associated with that context. We demonstrate that context-dependent generative traversal with CELTIC has the potential to reveal alterations in intracellular organization during context transitions.

## Results

### Deteriorated in silico labeling for rare cell populations

We used 3D spinning-disk microscopy images of genetically edited fluorescent human induced pluripotent stem cell lines (hiPSC) colonies from the Allen Institute for Cell Science (AICS) WTC-11 hiPSC Single-Cell Image Dataset v1 ^28^. The dataset comprises 3D field of view images, each containing label-free (brightfield), and a specific genetically edited EGFP-tagged protein representing an organelle. In addition, the dataset contains cell segmentation masks and metadata regarding the individual cells, that includes annotations regarding the cell cycle stage (interphase

/ prophase / early prometaphase / prometaphase-metaphase / anaphase-telophase-cytokinesis), and its location within a colony (interior / edge). We decided to focus on six organelles that span the range of in silico labeling performances reported in ^10^. Organelles with high performance in silico labeling included the nuclear envelope, actin filaments, and microtubules (average pixel-wise Pearson correlation coefficient of ∼0.78-0.88), organelles with intermediate performance included the mitochondria and endoplasmic reticulum (∼0.66-0.73), and organelles with low performance included the Golgi apparatus (0.23). We replicated the U-Net-based ^29^ in silico labeling model reported in ^10^ and reproduced their results (Table S1).

Our focus on single cell biologically meaningful context, required us to use the cells’ segmentation masks to isolate individual cells, and assign for each cell whether it was undergoing mitosis (non-dividing - interphase, or one of five mitotic stages), its location in the colony (whether it is located at the colony’s interior or edge), its volume, and its local density (i.e., number of adjacent neighboring cells in the colony). Overall we collected a single cell dataset consisting of 1,116-1,575 single cells per organelle, extracted from 100 fields of view images that were never seen by the trained model for each organelle. Cells in interphase accounted for more than 95% of the cells in our dataset, cells located away from the colony’s edge accounted for over 98%, cells with typical volumes (i.e., z-score higher than -1.5 relative to the overall population distribution) accounted about 93%, and cells in microenvironment of typical density (i.e., 5 or more adjacent neighbors) accounted for about 96% (Fig. 1B, Table S2). For each cell we measured the Pearson’s correlation coefficients (PCC) between the fluorescent ground truth and its corresponding in silico prediction. Overlaying a field of view PCC values onto the single cell segmentation masks showed poor endoplasmic reticulum in silico labeling for two cells in the prometaphase/metaphase stage of mitosis and for another cell located in a sparse neighborhood (Fig. 1C). Similarly, cells in prometaphase/metaphase and early prometaphase stages of mitosis showed poor in silico labeling for the nuclear envelope (Fig. 1D). To systematically evaluate this observation, we pooled all single cell PCC values across organelles and rare populations (Table S2). Cells undergoing mitosis displayed declined in silico labeling across all organelles compared to cells in interphase (Fig. 1E). The most dramatic performance deterioration occurred for microtubules and for the nuclear envelope. Cells located at the colony edge exhibited inferior in silico labeling of the microtubules and actin (Fig. S1A). Cells with small volumes and cells in sparse neighborhoods demonstrated reduced in silico labeling for most organelles, and most prominently for the nuclear envelope and actin (Fig. S1B-C). The decrease in silico prediction quality is a consequence of applying the model to under-represented data, likely due to out-of-distribution intracellular organization and the corresponding changes in these cells’ optical properties. We hypothesized that incorporating context information about each cell will increase the performance of in silico label models and address the generalizability issue of these models to under-represented data.

### CELTIC, cell-context dependent in silico labeling

We propose CELTIC, an in silico labeling model that integrates cellular contextual information. CELTIC is an image-to-image translation model, where the innermost layer of the network (“bottleneck layer”) explicitly encodes predefined cell context parameters. The single cell centric approach required us to move from a field of view-based to a single cell-based in silico labeling (Fig. 2A). Specifically, we cropped single cells according to their corresponding fluorescent plasma membrane-derived segmentation masks (see Methods). We defined five types of context representations per cell, that are briefly described here and are detailed in the Methods (Fig. 2B). The first type of context (“Stage”) was the mitotic state represented as a one-hot encoding vector. The second type (“Location”) was a binary indicator representing the cell’s location in the colony. The third context type (“Classic Shape”) captured the cell’s shape through a one-hot encoded vector derived from the clustering of shape descriptors (height, min/max width, volume). The fourth type (“Machine Learning Shape”) represented shape using autoencoder-compressed binary cell masks that were subsequently clustered, encoding each cell as a one-hot vector based on its shape cluster. Finally, the fifth context representation (“Neighborhood Density”) was a scalar quantifying the local neighborhood density measured as the number of adjacent cells. These five context representations were concatenated to define a 16-dimensional context feature vector encoding the single cell context.

**Figure 2.**
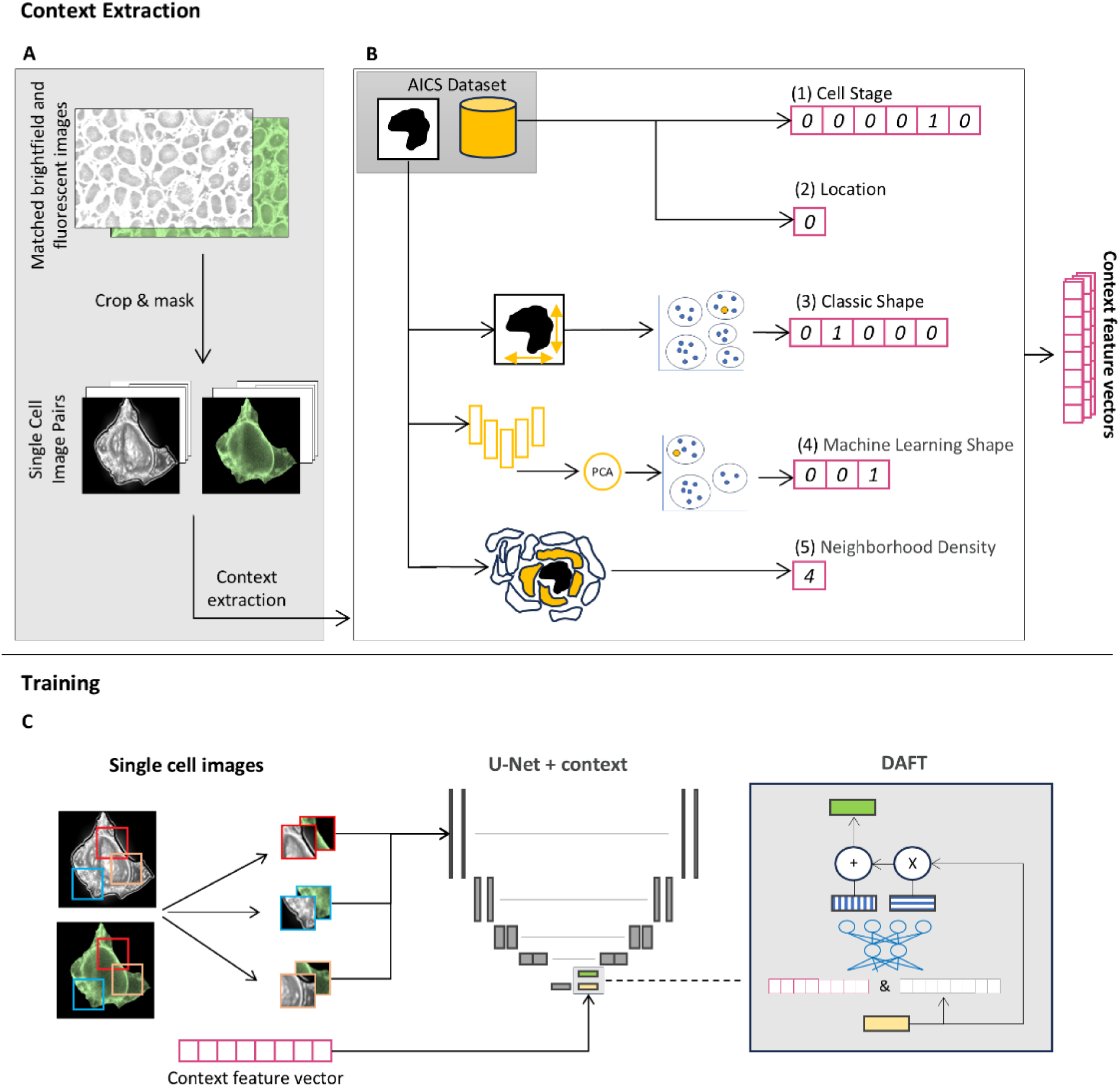
CELTIC - incorporating the cell context to the in silico labeling models. **(A-B)** Context extraction. **(A)** Single-cell images are cropped and masked. **(B)** Single cell context is extracted and represented by a 16-dimensional context feature vector, comprising five context types. The mitotic stage and the edge location indicator contexts were available in the Allen Institute of Cell Science (AICS) datatset’s metadata, other contexts are computationally extracted from the segmented cells and concatenated to define the context feature vector (magenta). **(C)** CELTIC’s architecture. Image patches are fed to CELTIC along with their corresponding context. The cell context is incorporated as an auxiliary input to the U-Net in silico labeling model transforming the U-Net’s bottleneck layer (yellow) into a context-enriched feature map (green) via the DAFT ^30^ block (gray box with a detailed view on the right). DAFT uses its own bottleneck to fuse the image and context, creating a scaler and shifter that adjust the feature map accordingly (see Methods and Fig. S2 for more details).

CELTIC extends the classic U-Net architecture by incorporating the context vector to its deepest layer. We followed the footsteps of a recent method for fusing image and tabular data called DAFT ^30^. DAFT was previously applied with ResNet for classification tasks and more recently with U-Net for medical image segmentation ^31^. DAFT uses the cellular context vectors to affine-transform the bottleneck image representations (Fig. 2C, Fig. S2). In CELTIC, this approach enables the network to learn a unified representation that incorporates both the intrinsic image details encoded at the network deepest layer along with the contextual cellular information.

### Cell-context contributes to the in silico labeling of rare cell populations

We compared CELTIC with the single cell U-NET to evaluate the contribution of the cell’s context to the in silico labeling predictions. We found that context contributed to better predictions of the endoplasmic reticulum and the nuclear envelope in dividing cells (Fig. 3A). Both models displayed poor performance in predicting microtubules mitotic spindle, but CELTIC was able to predict the two astral arrays radiating from the spindle poles during mitosis (Fig. 3A). These observations were corroborated by calculating the ΔPCC defined as the signed mean contribution of context to the single cell-based in silico labeling model, i.e., positive ΔPCC values indicate that the cell context contributed to the in silico prediction. Indeed, cell context enhanced the prediction of cells in mitosis for all organelles (Fig. 3B, Table S3). The most prominent contribution was measured for microtubules, which is expected given the dramatic changes occurring in microtubule organization during cell division. Similarly, inclusion of context improved the in silico labeling of cells at the colony edge, most notably for the Golgi apparatus and for microtubules (Fig. 3B). Cells with small volumes also showed improvement in most cases, particularly for the endoplasmic reticulum and for the nuclear envelope. Lastly, adding context enhanced the in silico prediction of cells in sparse neighborhood densities, with the most substantial differences observed for the Golgi apparatus and for the nuclear envelope (Fig. 3B). Intriguingly, context systematically contributed to the in silico labeling of most organelles, even when considering the entire cell population (Fig. 3B, “All”), but this contribution was marginal because the rare populations constituted only a small fraction of the dataset. It is important to emphasize that these models were trained without enrichment of rare populations. For instance only 28 of the 697 (4%) of the cells used to train the endoplasmic reticulum model were mitotic, with some subcategories, such as early prometaphase, containing as few as 2 cells, indicating that CELTIC learns meaningful representations from very limited data.

**Figure 3.**
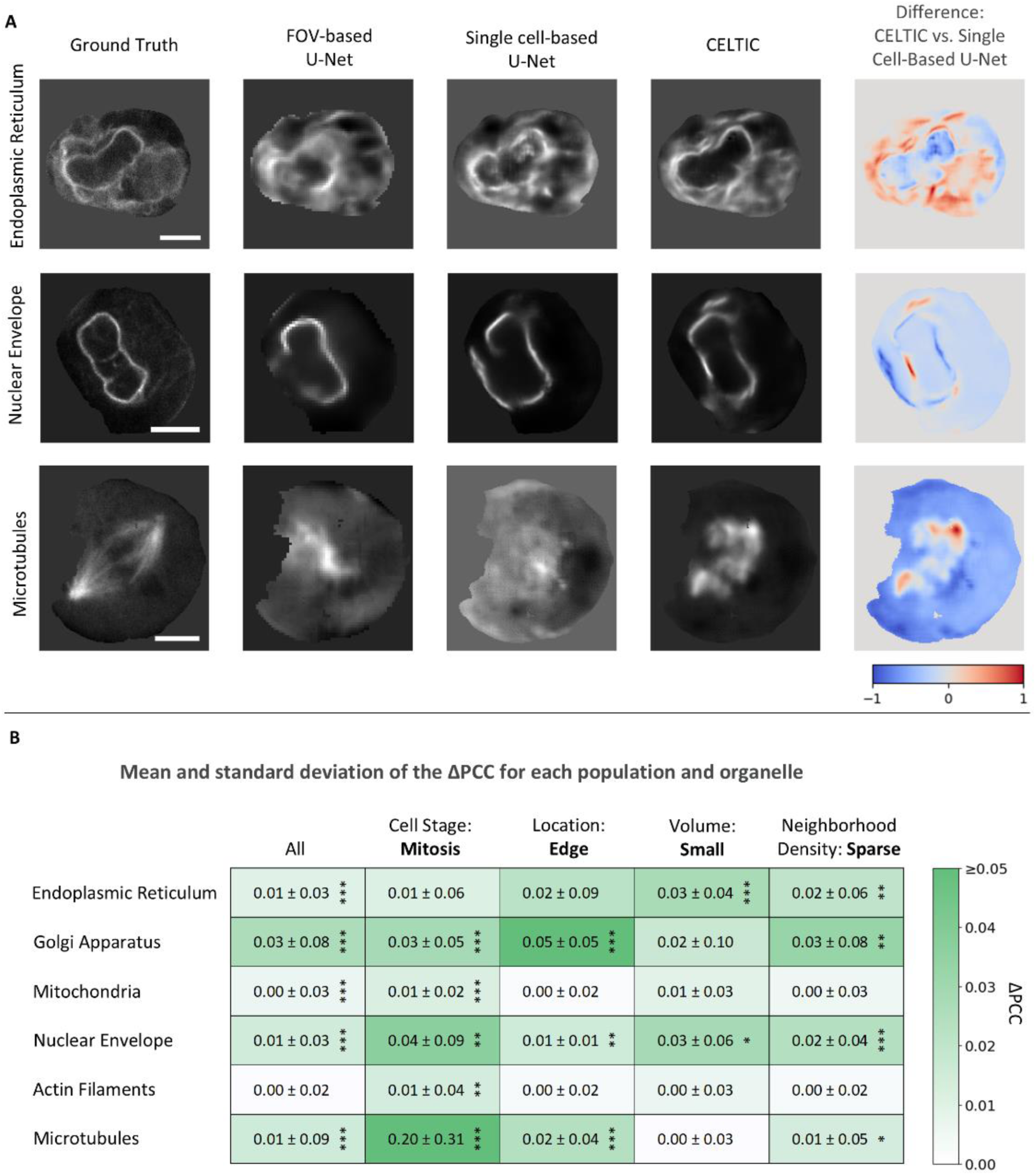
Qualitative and quantitative assessment of CELTIC’s contribution to the in silico labeling of rare cell populations. **(A)** In silico labeling visualization of mitotic cells. Each row visualizes a different organelle (top-to-bottom): endoplasmic reticulum of a cell in late mitosis, nuclear envelope of a cell in late mitosis, microtubules of a cell in the prometaphase/metaphase stage. Left-to-right: ground truth fluorescence, U-Net replicating ^10^, single cell-based U-Net, CELTIC, pixel-wise difference between the CELTIC model and the single-cell-based model. Red regions indicate positive intensity differences, while blue regions indicate negative intensity differences. The difference image is scaled from -1 to 1. Shown are Z slices that have been selected by an expert based on the ground truth full z-stack images. Scale bar = 5μm. **(B)** Quantification of the contribution of context, via CELTIC, to the in silico labeling of six organelles in the rare cell populations. Mean and standard deviation of the ΔPCC for each population and for each organelle. The entire population (left, “All”) for reference. Wilcoxon signed-rank test was used to reject the null hypothesis that ΔPCC is distributed around zero: * - p < 0.05, ** - p < 0.01, *** - p < 0.001. Full results are provided in Table S3.

To elucidate the contribution of each context type to the in silico labeling of rare populations, we conducted an ablation study. Specifically, we randomized each context type by shuffling the corresponding context values across the single cell population and measured the reduction in the in silico labeling with the shuffled context (Table S4). This analysis revealed that several context types were contributing to the in silico labeling of cells undergoing mitosis, unsurprisingly with cell stage ranking first among them (Fig. S3). The inclusion of the edge context predominantly impacted cells situated at the periphery of the colony for the in silico labeling of the endoplasmic reticulum, nuclear envelope and the microtubules. Cells with small volumes were primarily influenced by the shape contexts, particularly for the endoplasmic reticulum, Golgi apparatus, nuclear envelope, and actin. Intriguingly, the neighborhood density context had a minimal effect on most rare populations. Overall, we conclude that cell context, especially one that associates with a corresponding population, such as cell cycle stage for mitotic cells, contributes to in silico labeling by “guiding” context-aware representations that adapt to the different intracellular organizations associated with rare cell populations.

### Predicting spindle axis in mitotic cells enabled by CELTIC

The performance of an in silico labeling model should be evaluated based on its performance for specific application-appropriate downstream analyses, such as organelle localization, counts, and shape ^8,32,33^. During mitosis, the spindle axis connects the two centrosomes and is a critical determinant of cell division orientation and outcome ^34^. We decided to focus on the application-appropriate downstream analysis of determining the location and orientation of the spindle axis during mitosis from label-free images. We applied the single cell-based in silico model for microtubules, without and with the mitotics state context, for a set of prometaphase/metaphase cells that had not been seen before by the model. For consistency, we selected the middle z-slice from each image stack and resized the images to a standard size. Next, we performed a threshold-based segmentation of the in silico labeling predictions and set a line connecting the centers of mass of the two main contours in the image as the predicted spindle axis (see Methods). We defined two measurements to evaluate the predicted spindle axis: (1) location error - the distance between the center of the ground truth spindle axis line and the center of the predicted line (ΔC), and (2) orientation error - the angle between these two lines (Δθ) (Fig. 4A). Visual assessment indicated that while both models were not able to perfectly reconstruct the microtubules spindle axis, CELTIC was able to provide reasonable predictions regarding the spindle location and orientation (Fig. 4B). Quantification of ΔC and Δθ reported that the spindle axis was predicted by CELTIC with a median location error ΔC of 3.8 pixels (0.15-0.27 μm before resizing), and with a median orientation error Δθ of 13°. Comparison to the single cell-based model (without context) showed deterioration of over 2 fold in the predicted location (Fig. 4C) error and over 1.5 fold in the predicted orientation error (Fig. 4D). To confirm that these prediction errors were not achieved by chance we conducted a permutation test where we randomly shuffled all single cell predictions and calculated the median Δθ prediction errors across the shuffled population. This process was repeated 250,000 times and was used to calculate the statistical significance - the fraction of times that the permuted mean errors were smaller or equal to the observed (unpermuted) mean error, reaching statistical significance p-value < 0.00002 (Fig. 4E). These results suggest that the inclusion of cell context in CELTIC can enhance the in silico labeling of (application-appropriate) organelle localization patterns that are associated with that context.

**Figure 4.**
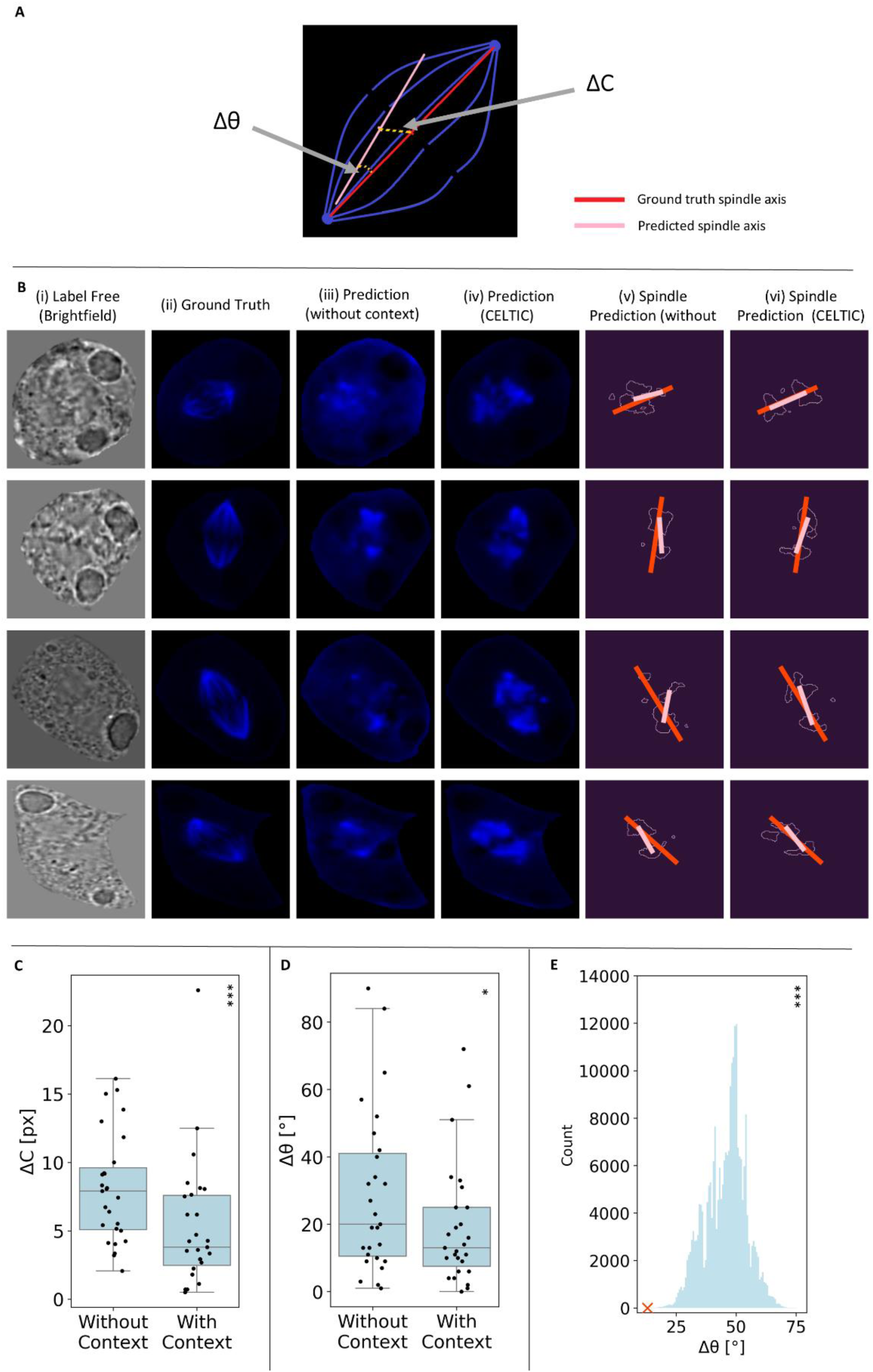
Application-appropriate downstream analysis: in silico prediction of the spindle axis location and orientation. **(A)** Measurements of the predicted spindle axis. The location error (ΔC) denotes the distance between the centers of the predicted (pink) and the ground truth (red) spindle axes. The orientation error (Δθ) denotes the angle between the predicted and the ground truth spindle axes. **(B)** Four representative cells (rows) in the prometaphase/metaphase mitotic state. Columns represent (left-to-right): (i) brightfield label-free image; (ii) ground truth fluorescent microtubules image from the z-stack’s middle slice; (iii-iv) single cell-based (iii) and CELTIC (iv) in silico labeling of microtubules; (v-vi) Threshold-based segmentation of the single cell-based (v) and CELTIC (vi) predictions, the spindle axis prediction (pink) and the ground truth (red). In silico labeling was performed in 3D, while spindle axis prediction and segmentation were conducted using a single 2D slice. **(C-D)** Distribution of the location (C) and orientation (D) errors of the single cell-based versus CELTIC prediction of the spindle axis. Each data point corresponds to a single cell. Wilcoxon signed-rank test was used to reject the null hypothesis that the values with CELTIC are similar or larger than with the single cell-based model. N = 27 cells, P-value (location) < 0.00017, P-value (orientation) < 0.028. **(E)** Permutation test: CELTIC’s observed orientation error (red ‘X’, median ΔC = 13°) and the histogram of random shuffling (N=250,000). P-value < 0.00002.

### Context-dependent generative organelle localization with CELTIC

The CELTIC representation incorporates the label-free image along with the cell context, which are jointly used to make the in silico prediction. This representation where the context can be manually manipulated can be used to generate a series of in silico labeling images of the same label-free image under varying contexts (Fig. 5A). Specifically, manipulating the context and assessing the corresponding alterations in the CELTIC-generated in silico labeling images to infer context-dependent changes in the intracellular organization. To illustrate the potential of this approach, we generated the integrated in silico labeling of the actin filaments, the nuclear envelope and the microtubules of a non-dividing cell where the actin filaments and the microtubules formed widespread networks throughout the cytoplasm, with a solid nuclear envelope surrounding the nucleus (Fig. 5B, top). Upon manipulating the cell to a prometaphase/metaphase mitotic context, for the same label-free image, CELTIC generated an integrated in silico labeling where (i) the nuclear envelope partially disassembled, (ii) the actin filaments are dispersed in the cytoplasm following the nuclear envelope disassembly and reorganized to form a ring at the cell equator in preparation for anaphase,, and (iii) the microtubules reorganized to form the aligned mitotic spindle (Fig. 5B, bottom). Repeating the same process in the opposite direction, we used the CELTIC generative organelle localization to transition a cell from prometaphase/metaphase to interphase. This process produced an integrated in silico labeling image resembling the interphase phenotype with dispersed actin filaments and microtubules in the cytoplasm, and a more rigid-appearing nuclear envelope (Fig. 5C). As another demonstration, we altered different contexts of the same cell. First, we manipulated the cell’s location context from interior to edge inducing altered localization of the actin filaments and microtubules toward the cell’s periphery (Fig. S4A, top versus middle), in concurrence with ^28^. Second, we manipulated the same cell’s stage context from interphase to mitotic (Fig. S4A, top versus bottom), demonstrating the generation of mitotic localizations, resembling the ground truth phenotypes (Fig. S4B). To systematically analyze how altering the cell’s context changes the corresponding CELTIC-generated image, we manipulated each of the five context types, and calculated the Pearson correlation score between the generated image before and after the context alteration for 230 cells from the dataset (Fig. 5D). Altering the cell stage context made the most dramatic change in the generated image space, particularly for the microtubules and the Golgi apparatus. Alteration of the location context was mapped to changes in the endoplasmic reticulum. Alteration of other contexts did not change much the corresponding generated images. Given the superiority of CELTIC on the corresponding cell populations (Fig. 3), these results showing context changes that do not translate to altered generated images can be explained by CELTIC’s generalized (pre-DAFT) representations, that better generalize to the corresponding rare cell populations. Altogether, the explicit representation of cell context enables guided traversal along the context axis of the same cell, overcoming the vast variability between different cells for easier visual interpretation ^35^. Analysis of the alteration in cell organization along these context traversals may teach us about the gradual change in integrated cell state along a physiological process.

**Figure 5.**
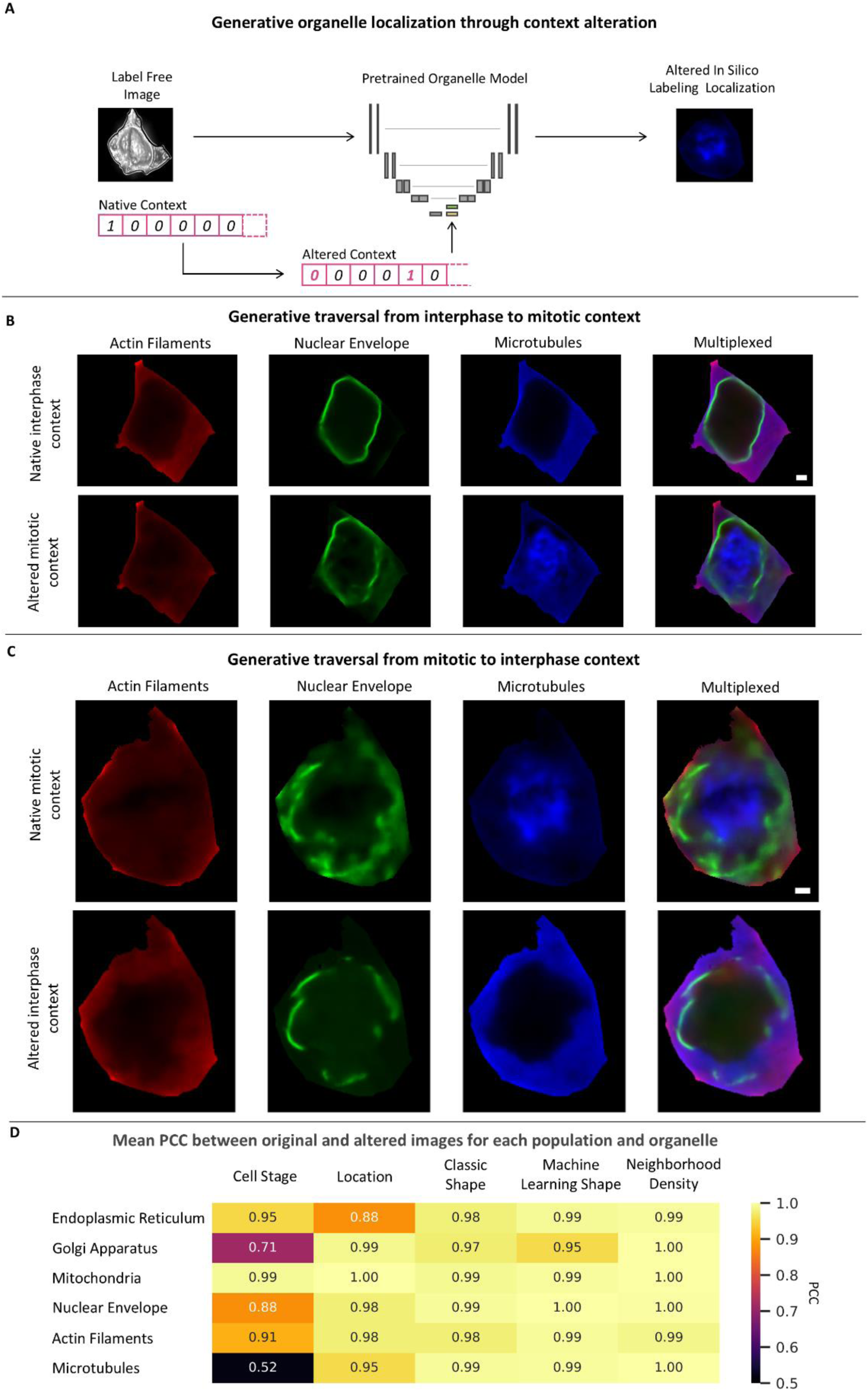
Context-dependent generative in silico labeling. **(A)** Approach: the native context feature vector is modified and used to generate an altered CELTIC in silico labeling. **(B)** In silico labeling of a cell in interphase. The native interphase context (top) versus mitotic context (bottom). Left-to-right: actin filaments (red), nuclear envelope (green), microtubules (blue), and a multiplexed representation of all three organelles together. Shown are the central Z slices. Scale bar = 2μm. **(C)** In silico labeling of a cell in prometaphase/metaphase. The native mitotic context (top) versus interphase context (bottom). The columns and scale bar are analogous to those in panel A. **(D)** Quantification of the changes in CELTIC-generated images resulting from context alterations. Six organelles, N = 230 cells. The Pearson Correlation Coefficient (PCC) was calculated between the original and altered images. Each cell in the table represents the mean PCC across the analyzed cells for the organelle (row) and the context type (column).

## Discussion

Measuring and interpreting how organelles adjust their internal structure and organization in respect to one another under different experimental conditions or during a physiological process is the “holy grail” of cell biology. In silico labeling is a promising method to overcome some of the technical hurdles that currently prevent us from reaching this ultimate goal. Here, we report that in silico labeling is confounded by rare cell contexts due to alterations in the cells optical properties that lead to out-of-distribution label-free images. CELTIC guides context-aware representations by incorporating the explicit cell context to the in silico labeling model. We show that CELTIC enhances the in silico labeling of rare cell populations, especially for organelle localization patterns associated with the corresponding context, and highlight its potential for modeling context-transitions through context-dependent generative capabilities. Our results emphasize the strong link between the cell’s context and its intracellular organization.

Cell context is a very broad term. It could be practically anything, from intrinsic cell contexts such as the ones shown in this study to extrinsic contexts such as the cell type, organelle, perturbations, disease state, assay, microscope, fluorescent marker and imaging parameters. The inclusion of context descriptors could be used to harmonize datasets from multiple resources to one large dataset. Thus context-dependent in silico labeling can be the enabler toward training general in silico labeling “foundation models”. Christiansen et al. made the first step in this direction by arguing for the benefit of training one model for the in silico labeling of multiple fluorescent channels ^12^, which in our case can be translated to providing the organelle type as context. An exciting possibility is to integrate the cell’s continuous state during the progression of a physiological process as context for in silico labeling. For example, using the FUCCI system as rich cell cycle context ^36^ or by computational prediction of the continuous cell state ^37–44^. Notably, the cell’s mitotic state, periphery location, and segmentation were available to us in the AICS dataset used in this study. In “the wild” these cell contexts would be computationally extracted from the raw images.

An alternative approach for incorporating context is through weakly supervised representation-learning, with the cell context as the “weak” label. Previous recent studies guided representations of protein localization patterns using the protein ^45,46^ or the perturbation ^47^ as the weakly supervised label. In principle, cell context can also be used as a label for weakly supervised representation learning. CELTIC’s explicit “injection” of the cell context to the representations has two major benefits. First, the possibility to control the context to morph a specific cell along the context trajectory, while fixing the other factors of cell-cell variability. This generative capacity along a context trajectory has the potential to serve as a hypothesis generation method to follow how the different organelles and their spatial inter-organelle dependencies are changing as a function of their contexts. The second benefit of explicitly injecting context over weak supervision is the exponential growth of the combinatorial context space with the number of different contexts. There are many possible contexts, and thus training representations to simultaneously encode multiple weak context labels is not feasible ^48^. CELTIC bypasses this limitation by avoiding the technical challenge of learning representations that encode context by providing it explicitly. Accordingly, the explicit context representation enables learning out-of-distribution label-free images from a very small set of examples consisting of tens of cells per rare population. One advantage of weak supervision over CELTIC is the simpler inference that does not require the weak context label as input.

Context does not improve in silico labeling for all organelles to the same extent. For example, in silico labeling of mitochondria was not improved much with the inclusion of context suggesting that the corresponding brightfield patterns used to localize the mitochondria do not depend much on the cell’s context. Systematic characterization of which contexts contribute to the in silico labeling of different cell populations can be used as a phenotypic signature of these populations, indicating how organelles reorganize upon physiological context alteration. Moreover, screening for the contexts that contribute most to the in silico labeling in CELTIC, compared to a model without context, can be used as a method to identify the cell’s context. This approach for context prediction is especially relevant for application-appropriate measurements, such as predicting the spindle axis, which should be dramatically improved during mitosis due to the strong association between intracellular organizational patterns and the context. Thus, the same approach can be used to discover unknown patterns that are associated with a specific cell context. Of course, this type of analysis must be carefully validated to rule out the possibilities that the pre-DAFT CELTIC representations generalized to the cellular context or that the model was not able to encode the organelle-specific context alteration due to lack of sufficient training data of rare populations.

## Methods

### Data

We used the Allen Institute for Cell Science WTC-11 hiPSC Single-Cell Image Dataset v1. From the field-of-view (FOV) spinning-disk confocal microscopy section, we used the 16-bit Z-stack images, acquired with a 100× objective, with a resolution of 624 × 924 pixels, and a physical pixel size of 0.108 x 0.108 µm. Each Z-stack consisted of 50–75 slices with 0.29 µm between consecutive slices. Specifically, we used the brightfield channel and the EGFP-tagged cellular structure channel, for the following proteins: alpha-tubulin (microtubules), beta-actin (actin filaments), lamin B1 (nuclear envelope), sec61B (endoplasmic reticulum), STGAL1 (Golgi apparatus), and Tom20 (mitochondria). For FOV segmentation, we used the cell segmentation channel, a 3D integer labelmap representing cell locations within the FOV. Lastly, we used the ‘edge_flag’ and ‘cell_stage’ features from the metadata CSV file.

We selected 80 FOV images per organelle to train and validate all models, that we call “Development set”. The images were selected from the list provided in the code repository of ^10^. In cases where images from this list were not available in the single cell dataset, we randomly replaced them with FOV images that closely matched in terms of their cellular properties, such as cell count, mitotic percentage, and edge characteristics. Additionally, we selected 100 FOV images for independent evaluations, based on the same criteria. We call these images the “Test set”. We used the FOV cell segmentation label maps to extract individual cells from the FOV images. We excluded cells that were not entirely inside the field of view due to the lack of metadata and inability to reliably extract shape descriptors. For each cell we constructed a 3D bounding box and applied the segmentation masks to the brightfield and EGFP FOV images, resulting in a two channel single-cell dataset. Pixel intensities of all images were normalized to have a mean of 0 and a standard deviation of 1 to account for variations in illumination intensity.

We categorized each cells into populations based on the following criteria: Stage (mitosis vs. interphase) by using the metadata cell_stage field; Location (edge vs. interior) by using the metadata edge_flag field; and Volume, where small cells were those with a volume lower than - 1.5 standard deviations from the mean. For neighborhood density analysis, cells were considered in sparse neighborhoods if they had fewer than 5 adjacent cells.

### In silico labeling replication

To replicate the in silico U-Net model in ^10^ we used the Development set. For training we used 64 FOV images (56 for training, 8 for validation), and for testing we used the remaining 16 FOVs. We followed the preprocessing steps that were reported in the paper: resizing the z-slices to 244 × 366 pixels and normalizing each image to a mean of 0 and a standard deviation of 1. We refer to these models as ‘FOV-based models’ and report their replication results in Table S1.

We used the Test set to assess the in silico labeling performance of the FOV-based models at the single-cell level. The procedure for extracting individual cells was adjusted (see Methods/Data). Specifically, we resized the segmentation channel to match the size of the FOV images, without smoothing, to ensure consistency of the segmentation at the resolution of a single pixel. We then applied a binary mask to both the predictions and their corresponding fluorescent targets and calculated the Pearson correlation coefficients (PCC) between them, considering only the pixels belonging to the cell, as defined by the segmentation mask.

To assess the null hypothesis that the distributions of the rare and non-rare populations are identical, with no significant difference in their medians, we employed the Mann-Whitney U test (results in Table S2).

### Context Representations

#### Stage

The representation was extracted from the single-cell metadata CSV file, where the cell cycle stage is indicated as one of six stages: “M0” (interphase), “M1M2” (prophase), “M3” (early prometaphase), “M4M5” (prometaphase/metaphase), “M6M7_single,” and “M6M7_complete” (anaphase/telophase/cytokinesis in two stages). This annotation was generated by a deep learning-based classifier and rule-based criteria ^28^. We represented this information numerically as a six-column one-hot vector, with each column corresponding to one of the six cell cycle stages.

#### Location in the colony

The representation was extracted from the edge_flag column in the single-cell metadata CSV file (’true’ for edge cells, ‘false’ for interior cells). We represented this context as a boolean.

#### Shape representations

We used two types of shape representations. The first shape representation was of predefined shape features capturing classic shape attributes such as cell volume termed “classic shape”. The second shape representation was machine-learning derived shape features that captured more subtle and complex attributes, which are not necessarily encoded in the “classic” representation, and termed “machine learning shape”. These two representations are described next.

#### Classic Shape

We used the segmentation masks to calculate the cell’s height (defined by the minimal and maximal z coordinates within the mask), cell volume (the volume enclosed by the mask), and cell width (the range of sizes along the x and y axes, combined into a single xy axis). Following min-max scaling of these measurements, we applied a k-means clustering with k=5, as determined by evaluating the relationship between the number of clusters and the within-cluster sum of squares. Each cell was assigned to a cluster, which was represented as a one-hot encoding vector. We also attempted at incorporating the distances of each cell to the k-means centroids but found this to be less effective.

#### Machine Learning Shape

We resized each cell’s (binary) segmentation mask to an image of size [32, 64, 64] for z, y, and x, respectively. We trained an autoencoder to encode representations of these single cell segmentations. The encoder consisted of two 3D convolution layers (depths 16 and 32), each followed by ReLU activation and max-pooling with kernel size 2 and stride 2. The decoder included two 3D transposed convolution layers, with ReLU activation for the first and sigmoid activation for the last. The autoencoder was trained for 10 epochs to minimize the mean squared error (MSE), on the cell segmentation images from the training set. The latent space of the autoencoder, shaped as [32, 8, 16, 16], was reshaped into vectors of size 65,536. Principal Component Analysis reduced the dimensionality to 5 main components. K-means clustering was then performed, determining the optimal number of clusters (k=3). Shape clusters were represented by cluster membership, with a binary indicator for cluster belongingness.

#### Neighborhood Density

For each cell we identified and counted the neighboring cells according to the segmentation image of the field of view. The number of neighboring cells was min-max scaled according to the minimum and maximum number of neighbors in the training set.

### CELTIC architecture

We adopted the 4-level U-Net architecture from ^10^, incorporating convolutional layers, batch normalization, and ReLU activations. The complete architecture is detailed in Fig. S2. Similarly to the original model, which was trained on field-of-view images, we employed patching, as even a single cell in 3D was too large to fit into the GPU. We used random patches of size 32 × 64 × 64 (z, y, x), excluding those that contained no signal within the segmentation mask. We integrated the DAFT ^30^ to the U-Net’s architecture. DAFT extends the FiLM ^49^ method to fuse image data with its complementary tabular information. Integrating image and tabular data is challenging due to their dimensionality mismatch, and naive approaches like concatenating latent representations often result in networks prioritizing image features, with minimal improvements over traditional CNNs ^49,50^. The DAFT method addresses this limitation by facilitating a dynamic exchange of information between the 3D image and the tabular data through an auxiliary neural network, enhancing their interaction and integration capacity. Specifically the U-net’s bottleneck produced a feature map tensor (of size 512 × 2 × 4 × 4) that was provided as input to the DAFT architecture. This input underwent 3D adaptive average pooling, resulting in a single value per feature map. This tensor was concatenated with the 16-dimensional context feature vector and encoded by a fully connected layer with a compression rate factor, which was a hyperparameter, followed by a ReLU activation function. Several bottlenecks were tested (2, 4, 12, 32, 48, 128), and the final chosen factor for each organelle is reported in Table 3. A subsequent linear layer decoded the representation to twice the original feature map size. Half of this decoded representation was activated by a sigmoid function and used to scale the feature map values, while the other half shifted them. The adjusted feature map was then propagated through the U-Net’s upstream layers. The alternative configurations of applying DAFT to different or additional layers showed inferior results and thus were not further pursued.

### Models’ training and performance evaluation

To evaluate the contribution of cell context, we trained and evaluated in silico models for each organelle dataset: CELTIC (with the DAFT block and context) versus U-NET. We randomly split the Development set into three subsets: “train”, “validation 1”, and “validation 2” with a ratio of 7:1:2. The split was performed at the FOV level (rather than the single cell level) to ensure that cells from the same FOV appeared in only one data subset, thereby preventing the models from learning batch effects. The models were trained for 60,000 iterations, using mean squared error (MSE) as the loss function, calculated on the masked signal area. During training, MSE was evaluated on the “validation 1” subset once every 100 iterations, and the best model was selected based on the iteration that achieved the lowest validation score. To determine the optimal bottleneck size hyperparameter for DAFT, we trained both the U-NET and the CELTIC models using three different split seeds of the Development set. We selected the bottleneck size that maximized the CELTIC performance on the “validation 1” subsets. From the three models of the best bottleneck, we selected the one that performed best on rare populations in the “validation 2” subsets. This process resulted in a total of 12 models, a matched CELTIC and U-NET models for each one of six organelles. All training was conducted on an NVIDIA RTX 2080 GPU using PyTorch.

These six pairs of matched organelle-specific CELTIC-versus-U-NET models were evaluated using the Test set. For each cell we calculated the in silico labeling PCC for CELTIC versus the matched U-NET prediction. We evaluated the contribution of context via CELTIC for five cell populations: one population containing all cells (“All”), and four rare populations (mitosis, edge, small volume, sparse neighborhood density ). For each population we calculated the ΔPCC = CELTIC_PCC_ - U-NET_PCC_ and the corresponding Wilcoxon signed-rank test to reject the null hypothesis that the ΔPCC values are distributed around zero. The p-values, population sizes and distribution of PCC values across organelles and rare populations are detailed in Table S3.

### Ablation study

To systematically assess the contribution of context to the in silico labeling of rare cell populations, we performed bootstrapping analysis by permuting the context between cells and evaluating the resulting deterioration in the in silico labeling performance. Specifically, for each organelle, context type, and rare population, we shuffled the values of the context type across all cells. We then used the pre-trained model to generate predictions on the Test set images with permuted context and calculated the mean PCC across shuffles.This process was repeated 10 times for each representation using different random seeds. The results, reported in Table S4 and in Fig. S3, were compared to the mean PCC obtained with true (unshuffled) context. Statistical significance was assessed using paired t-tests between the true and shuffled contexts.

### Application-appropriate assessment of the predicted spindle axis

As an application-appropriate assessment we evaluated the ability to determine the location and orientation of the spindle axis during mitosis from brightfield images. We used all 27 cells in the mitotic prometaphase/metaphase stage from the microtubules Test set data. We performed in silico labeling of microtubules using the corresponding CELTIC and U-NET model. To segment these in silico predictions and calculate the spindle axis, we devised the following pipeline. First, resize the middle z slice of the 3D images to 128×128 pixels. Second, erode the microtubules prediction according to the cell’s segmentation mask with a kernel size of 5 pixels to remove any residual predictions on the cell’s border. Third, threshold the microtubules prediction according to the 90th pixel intensity percentile. Forth, to define the spindle axis, we selected the two largest connected regions in the segmented image, calculated their moments and their center of mass, and defined the spindle axis as the line connecting the two centers of mass. In cells with a single region we defined the spindle axis according to the longest chord of the region, i.e., according to the longest line that could be accommodated within this region. To define the ground truth spindle axis we manually annotated the 27 cells. The location error (ΔC) was calculated as the distance between the predicted and the ground truth spindle axes. The orientation error (Δθ) was calculated as the cosine of the angle between the predicted and the ground truth spindle axes. Statistical significance was determined with a permutation test, in respect to the Δθ achieved in 250,000 random shuffles of the ground truth values.

### Using context for image generation

To generate how an organelle may change for the same cell in a different context, we used CELTIC to predict the in silico labeling of the same cell with different context inputs. For example, to generate the images in Fig. 5A, we in silico labeled an interphase cell with the actin filaments, nuclear envelope and microtubules CELTIC models. The top image is a result of the prediction with the native interphase context. For the bottom result, we manipulated the cell stage representation in the context vector from interphase (mitotic context representation of ‘100000’) to prometaphase/metaphase (‘000100’). To quantify how altering context affects the in silico labeling we used 230 cells from 16 unseen FOVs. For each cell and for each organelle we used CELTIC with the appropriate context, and then repeated the inference by manipulating the context vector. We measured the Pearson correlation coefficient between these context-altered in silico labeled predictions.

## Code and data availability

We are currently organizing our source code and processed data. We will make both publicly available at https://github.com/zaritskylab/CELTIC as soon as possible (before journal publications).

## Funding and Acknowledgments

This research was supported by the Israel Science Foundation (ISF, grant No. 2516/21, to A.Z), and by the Israeli Council for Higher Education (CHE) via the Data Science Research Center, Ben-Gurion University of the Negev, Israel (to AZ). We thank Matheus Viana, Nathalie Gaudreault, Jianxu Chen, Sussane Rafelski, Michal Feldman and Natalie Elia for discussions, suggestions and critical reading of the manuscript.

## Author Contribution

AZ and NE conceived the study. NE developed the computational method and analyzed the data. NE and AZ interpreted the data and drafted and edited the manuscript and approved its content. AZ mentored NE.

## Supplementary figures

**Figure S1.**
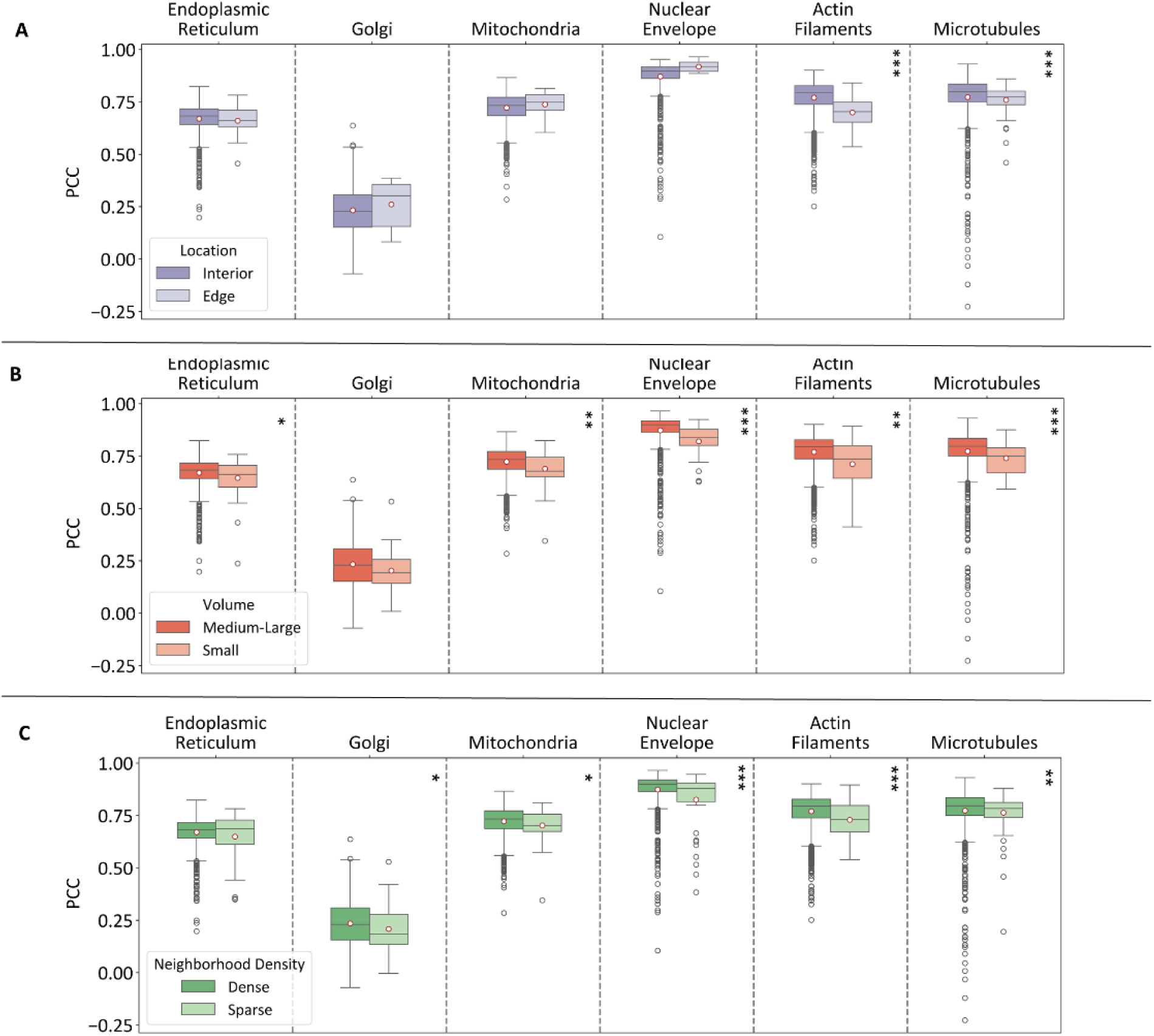
Distribution of single cell in silico labeling performance across organelles for rare cell populations. **(A)** Location within the colony (interior versus edge), (**B**) Volume (Medium-Large vs. Small). **(C)** Neighborhood density (sparse vs. dense). Mann-Whitney U test: * - p < 0.05, ** - p < 0.01, *** - p < 0.001. Full results are provided in Table S2.

**Figure S2.**
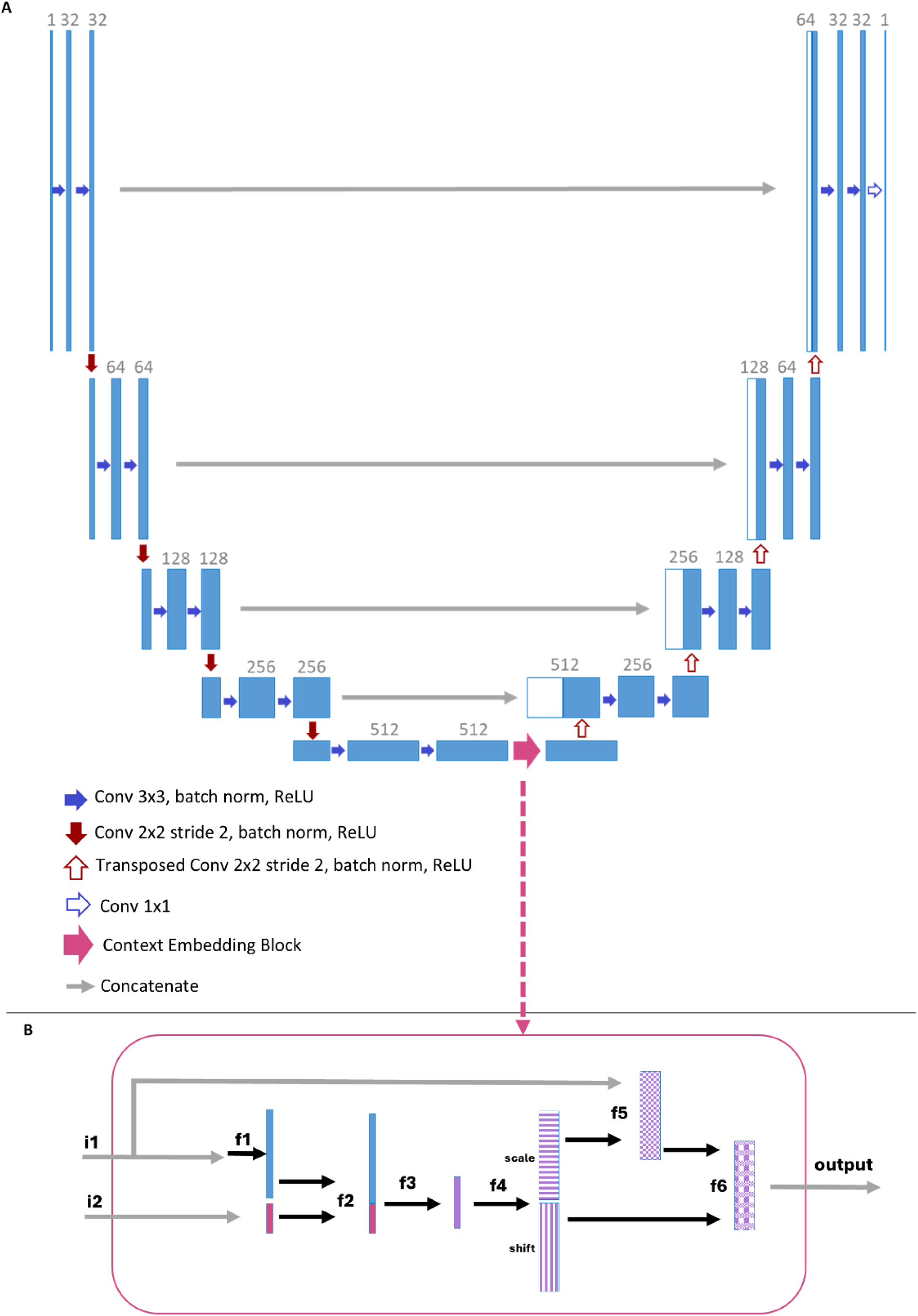
The CELTIC architecture. (**A**) The U-Net architecture with the CELTIC context embedding block (magenta arrow) adapted from Ronneberger et al. ^29^. (**B**) Context Embedding Block operation (magenta rectangle) adapted from ^30^, where i1 corresponds to the bottleneck feature maps from the U-Net, and i2 represents the context vector as input. Black arrows illustrate the flow between internal components within the block. The actions are denoted as follows: f1 - global average pooling; f2 - concatenation; f3 - fully connected layer with ReLU activation; f4 - fully connected layer with post separation for scaling and shifting; f5 - multiplication of i1 by the scale, followed by sigmoid activation; f6 - addition of the shift to the result of f5.

**Figure S3.**
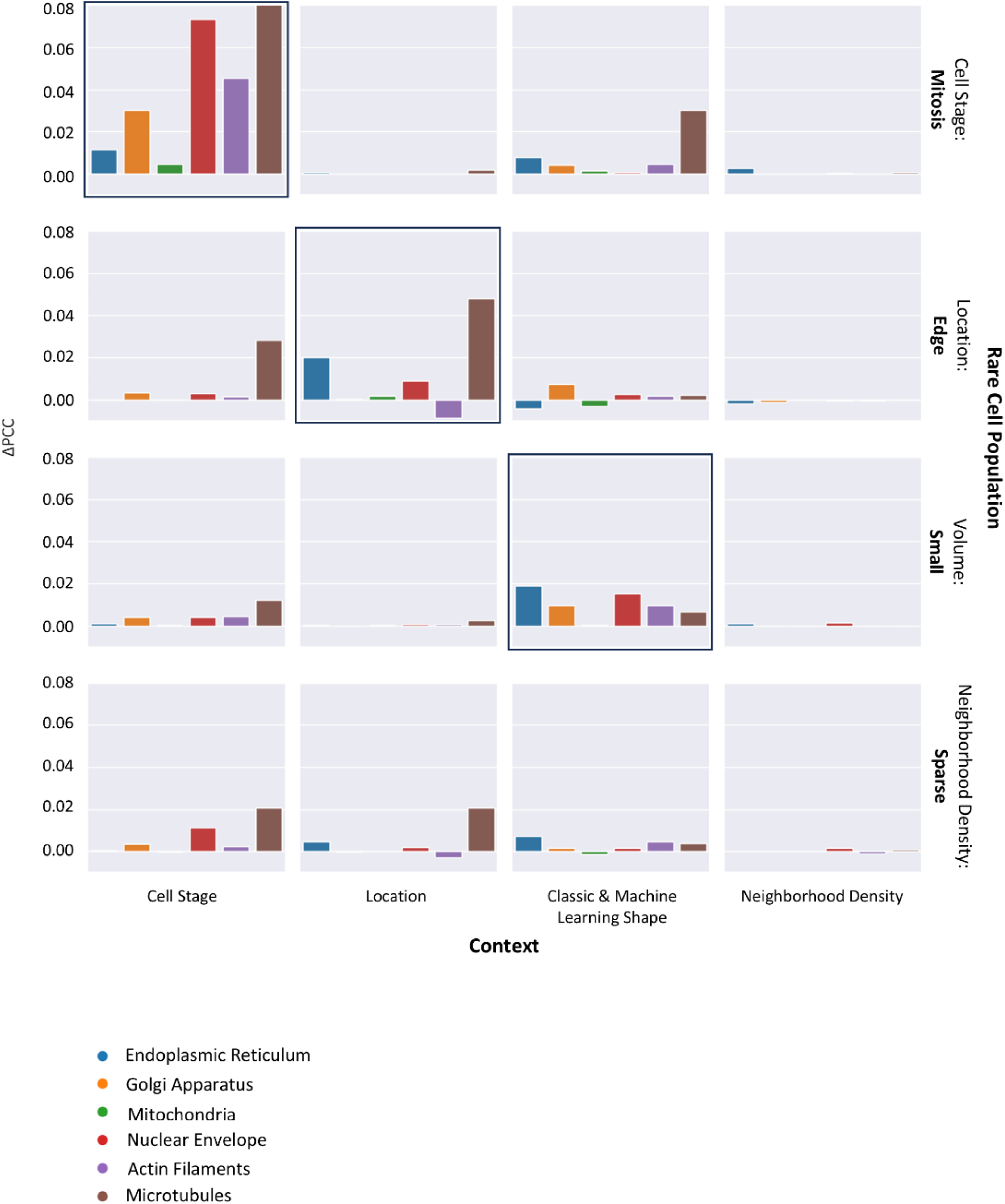
Ablation study assessing the contribution of the different context types to the in silico labeling of rare cell populations. Each row represents a rare cell population, and each column represents a shuffled context type. The bar plots depict the mean Pearson correlation coefficient difference (ΔPCC) between shuffled and non-shuffled contexts for each organelle (colorcoded, see legend). Black square borders indicate that the rare population (row) is most affected by its corresponding CELTIC cell context type, shown for mitosis, edge, and small volumes.

**Figure S4.**
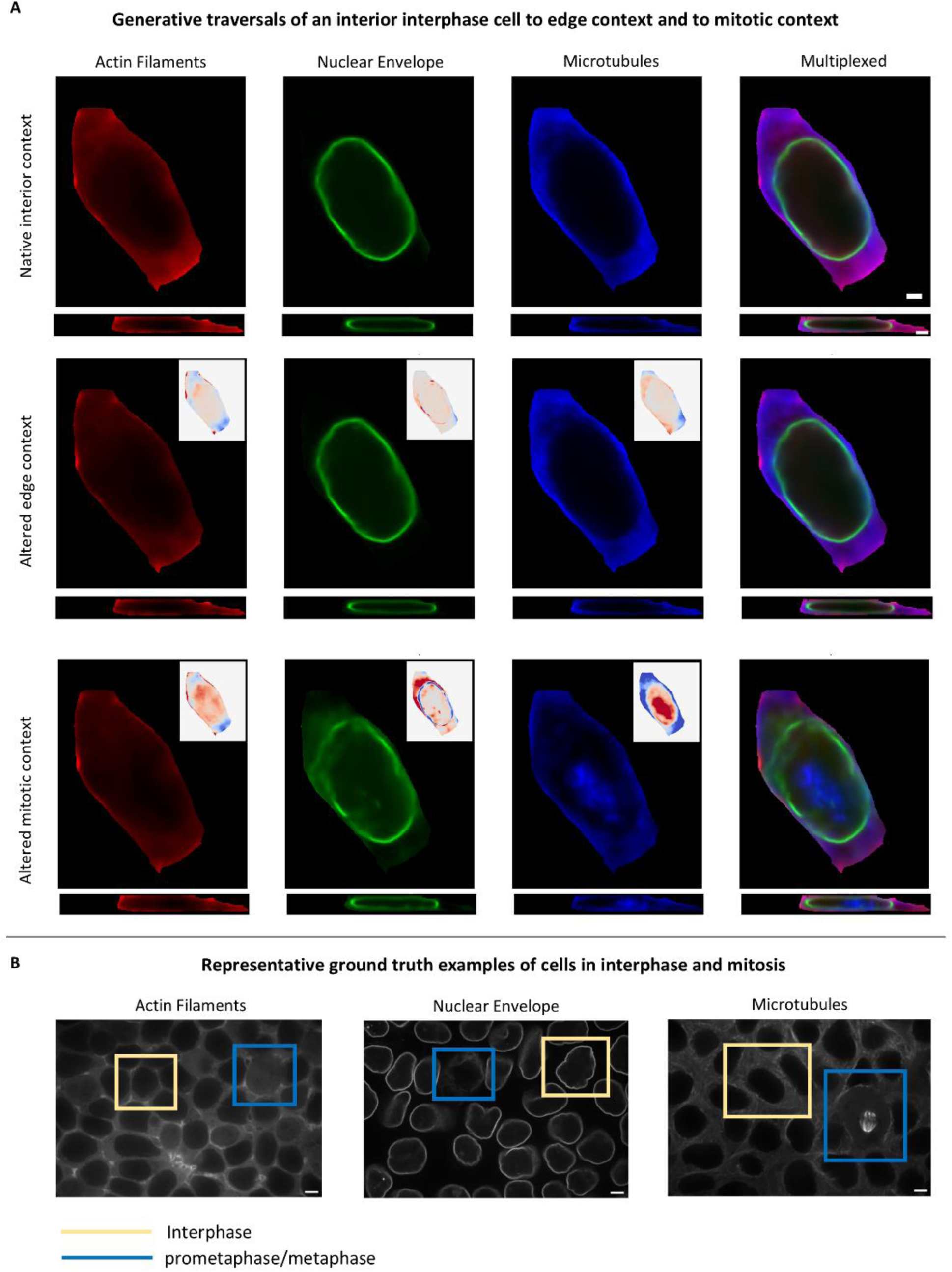
Context altered in silico labeling of the same cell. (**A)** CELTIC in silico labeling of a cell in interphase, located at the interior of the colony under different context alterations. Top-to-bottom: native interior context, colony edge context, mitotic context. Left to right: actin filaments (red), nuclear envelope (green), microtubules (blue), and a multiplexed representation of all three organelles together. Shown are XY and their corresponding YZ views in the central Z/X image spaces correspondingly. Scale bar = 2μm. Inset heatmaps (top-right) indicate the difference between each altered-context generated image and its corresponding native-context generated images, scaled between -1 (blue) and 1 (red). (**B**) Ground truth field of view images of the organelles and cell stages as generated in Fig. 5B-C and Fig. S4A (top and bottom rows). Yellow squares highlight an example of a cell in interphase, blue squares highlight a cell in prometaphase/metaphase. Scale bar = 5 μm.

